# From seed to flour: sowing sustainability in the use of cantaloupe melon residue (*Cucumis melo* L. var. *reticulatus*)

**DOI:** 10.1101/676916

**Authors:** Josiane Araújo da Cunha, Roseane Claro Nabas, Priscilla Moura Rolim, Karla Suzanne Florentino da Silva Chaves Damasceno, Francisco Canindé de Sousa Júnior, Larissa Mont’Alverne Jucá Seabra

## Abstract

Reduction of waste from food industry and food services is a current concern due to the large amount of waste generated, including peels and fruit seeds. The objective of this study was to obtain a flour produced from Cantaloupe melon seeds (*Cucumis melo* L. var. *reticulatus*) and to evaluate the viability of using the product as an ingredient in cake manufacturing. In this study, different formulations were developed: standard cake – 0% (F1) and cakes containing melon seed flour as substitute of wheat flour in 10% (F2), 30% (F3), and 50% (F4) concentrations. Centesimal composition, dietary fibre, structural and morphological characterization, determination of mineral composition, and evaluation of fatty acids profile in melon seed flour were carried out. To determine the overall acceptance of cake formulations, sensory analysis was performed with 135 non-trained panelists, which also included the identification of sensorial attributes using the *Just About Right* ideal scale test. The results showed that the melon seed flour has considerable nutritional value, with 18% proteins, 3% moisture, 4% ash, 30% lipids, and 35% dietary fibre. Melon flour also has a significantly high content of minerals, mainly phosphorus (1507.62 mg/100 g), potassium (957.35 mg/100 g), and magnesium (504.03 mg/100 g). The polyunsaturated fatty acid fraction was the most abundant in melon seed flour, with predominance of omega-6 fatty acids (17.95 g/mg of sample). Sensorial analysis disclosed good acceptance for formulations containing 10% and 30% of melon seed flour, with the 10% formulation being the most accepted. The research showed the feasibility of using the melon seed flour in cake production, as well as the possibility of using food waste in restaurants and food industries in order to adhere to sustainable production actions.

## 1. INTRODUCTION

The wastage of natural resources has been a great concern worldwide needing the adoption of new behaviours regarding their utilisation. In this context, awareness of the individuals regarding the control of waste generation is of great importance and involves prevention practices and the integral use of food. Sustainability, especially the use of wasted edible parts, is an essential component for minimising negative impacts on the environment [1].

Conscious production has become one of the major challenges to be addressed by the food industry, whose main concerns are cost reduction and increasing customer satisfaction and needs [2]. Reducing food waste has positive impacts, both economically and environmentally, and is essential for a sustainable food system to guarantee food and nutritional security. It is therefore necessary to understand the reasons for the wastage, as well as to implement measures for identification and prevention through the monitoring of the entire production process [3].

To protect the environment, the United Nations [4] adopted a sustainable development agenda to be implemented by 2030. When analysing the 17 proposed objectives, objective 12 establishes a guideline “to ensure sustainable production and consumption patterns”, aiming at the efficient use of natural resources, reduction of food waste and waste generation, encouraging the adoption of sustainable practices, and ensuring that people are made aware of sustainable development.

The absolute need to preserve the environment has resulted in reconsideration of the practices of waste minimisation, including those generated in foodservices. The practice of environmental preservation comprises a set of policies and strategies, aimed at reducing and controlling the resultant impacts [5]. Encouraging full utilisation of food, using for example, fruit seeds and peels offered foodservices, contributes to the achievement of sustainable production in meals. Seeds from various foods have been used in order to provide new products, and have becoming alternative food sources, as well as excellent natural sources of nutrients [6].

Melon consumption is very high worldwide, particularly in European countries. The Cantaloupe melon *(Cucumis melo* L. var. *reticulatus)*, largely present in Brazil, is typically regional. The state of Rio Grande do Norte has leadership in area and production [7]. Due to its pleasant flavour and aroma, the fruit appears in the menus of most foodservices and generates a high volume of residues from its peels and seeds, which are discarded because they are not usually edible. Throughout the melon production and consumption chain, the utilization ranges from 38 to 42%, that is, 58 to 62% of the raw material is discarded as waste [8]. Underutilised food sources like these have the potential to contribute substantially to food and nutrition security, and to improve ecosystem functions [9].

Melon seeds contain high percentages of lipids, proteins, and fibres. They present antiproliferative activity and antioxidant properties, in addition to their prebiotic potential [10,11]. In addition, the presence of flavonoids and phenolic compounds reaffirms that the melon seed flour *Cucumis melo L*. has high antioxidant properties [12]. They contain significant amounts of minerals such as magnesium, phosphorus, sodium, and potassium [13]. Melon seeds are rich in lipids, mainly polyunsaturated fatty acids, especially linoleic and linolenic [14,15]. Additionally, they have considerable amounts of essential amino acids, such as isoleucine, methionine, tyrosine, phenylalanine, and valine [16].

The foodservices can add to their objectives values such as the sustainability guarantee and good acceptance of new preparations. The use of parts conventionally not consumed, such as melon seeds, in the development of a product that is part of the habitual diet, is a good option to the reduction of waste, as well as a strategy for the improvement in foodservices, through the implementation of sustainable menus [17]. Bakery products, like cake, becomes an alternative to the addition of new ingredients such as flours of fruit waste, conditioning it to the expectation of good sensory acceptance [18]. Thus, this research aimed to obtain a flour from the Cantaloupe melon seeds, to characterize it and to evaluate the viability of its use as an ingredient in the development of cakes in institutional foodservice.

## 2. METHODOLOGY

### 2.1 Melon Seed Flour

Cantaloupe melon seeds of the *reticulatus* variety, commercially known as Japanese melon, were collected on the same day of distribution of the fruit to the consumers (April 2018), in the University Restaurant of the central campus of Federal University of Rio Grande do Norte, Brazil. Approximately 5.0 kg of melon seeds were transported in an isothermal box to the Dietetic Laboratory.

The seeds were weighed, washed in tap water to eliminate the residual pulp, and sanitised with sodium hypochlorite solution (200 ppm) for 15 min. Sequentially, they were submitted to a technological drying process using a ventilated oven (TECNAL brand, model T6394/2) at 80°C for 24 h [19]. A domestic 150W grinder (Cadence brand) was used to obtain the Melon Seed Flour (MSF). MSF was kept at room temperature and protected from light until analysis.

### 2.2 Centesimal Composition

Centesimal composition and dietary fibre analyses followed the methods used by the Adolfo Lutz Institute [20]. Moisture was determined by direct drying in an oven at 105°C until constant weight. Ashes were obtained by heating and incinerating the sample at 550°C (muffle). Continuous extraction of the lipids was carried out in a Soxhlet type apparatus, followed by evaporation removal of the solvent used (ether). Protein determination was based on the Kjeldahl nitrogen definition, and the obtained value was multiplied by a factor of 6.25, transforming the number of g of nitrogen into number of g of proteins. The dietary fibre was established by the enzymatic-gravimetric method, which is based on the quantification of the total food fraction after enzymatic digestion [21. All analyses were performed in triplicate, with the exception of the total dietary fibre analysis.

### 2.3 Mineral Analysis

Mineral analysis was carried out by optical emission spectrometric induction with inductively coupled plasma – ICP OES. The sample of melon seed flour was incinerated in the Bromatology Laboratory of UFRN to obtain ashes. The material was diluted with 50 mL of nitric acid and then assigned to the Nucleus of Primary Processing and Reuse of Produced Water and Waste – NUPPRAR, for analysis of the microelements: calcium, copper, total chromium, iron, total phosphorus, magnesium, manganese, total nickel, potassium, selenium, sodium, and total zinc.

### 2.4 Fatty acids profile

The composition of fatty acids was determined in the Food Technology Institute – ITAL. The saponification process was carried out with 2% NaOH solution in methanol, followed by esterification with ammonium chloride solution, and sulphuric acid in methanol. The methyl esters of the prepared fatty acids were quantified by gas chromatography. Quantification was performed by area normalisation and the results for each fatty acid were expressed in g/100 g of sample [22, 23].

### 2.5 Morphological Analysis

The microstructure analysis was performed using Scanning Electron Microscopy (SEM), Hitachi Tabletop Microscope TM-3000 model. A small amount of flour was fixed with carbon tape in metal support *(stubs)* and metallised with a thin layer of gold *(sputtering)* to be submitted to vacuum under a voltage equivalent to 30 mA/min. The sample was then fixed in the sample holder for the SEM analysis and positioned in the apparatus for image projection. The resulting images were evaluated by secondary electrons, accelerated at a 15-kV voltage and captured in digital form, with increasing voltages varying from x200 k to x4.0 k. Microscopic analysis, using a bench Magnifying Glass (Oleman), aiming at the size of the flour granules was also carried out.

### 2.6 Sensory Analysis

#### 2.6.1 Panelists

Sensory evaluation was conducted in the Sensory Analysis Laboratory of UFRN Nutrition Department by a team of 135 non-trained panelists. The testing included the Analysis of Global Acceptance (nine-point scale) and *Just About Right* (JAR), a method utilised to determine the presence of some attribute(s) and/or intensities, in this case, flavour (sweet) and texture (softness) [24,25]. As inclusion criterion, the individuals who constituted the university community (18 to 60 years old) were selected to take part of the sensory analysis, and the individuals who claimed oral injury, allergy, or intolerance to the used ingredients were excluded.

#### 2.6.2 Cake Formulations

Melon seed flour was used as raw material for the cakes, with the simple cake standard formulation being the basis for the development of the others. Therefore, in addition to the traditionally prepared cake at University Restaurant, three other products were prepared, with different contents of MSF, partially replacing the wheat flour, in the Laboratory of Dietary Techniques of UFRN Nutrition Department.

Table 1 presents MSF cake formulations, related to wheat flour content: formulation 1 (F1: 0% MSF and 100% wheat flour); formulation 2 (F2: 10% MSF and 90% wheat flour); formulation 3 (F3: 30% MSF and 70% wheat flour); and formulation 4 (F4: 50% MSF and 50% wheat flour). The proportions of other ingredients (sugar, chemical yeast, milk, margarine, and eggs) remained unchanged.

**Table 1.**
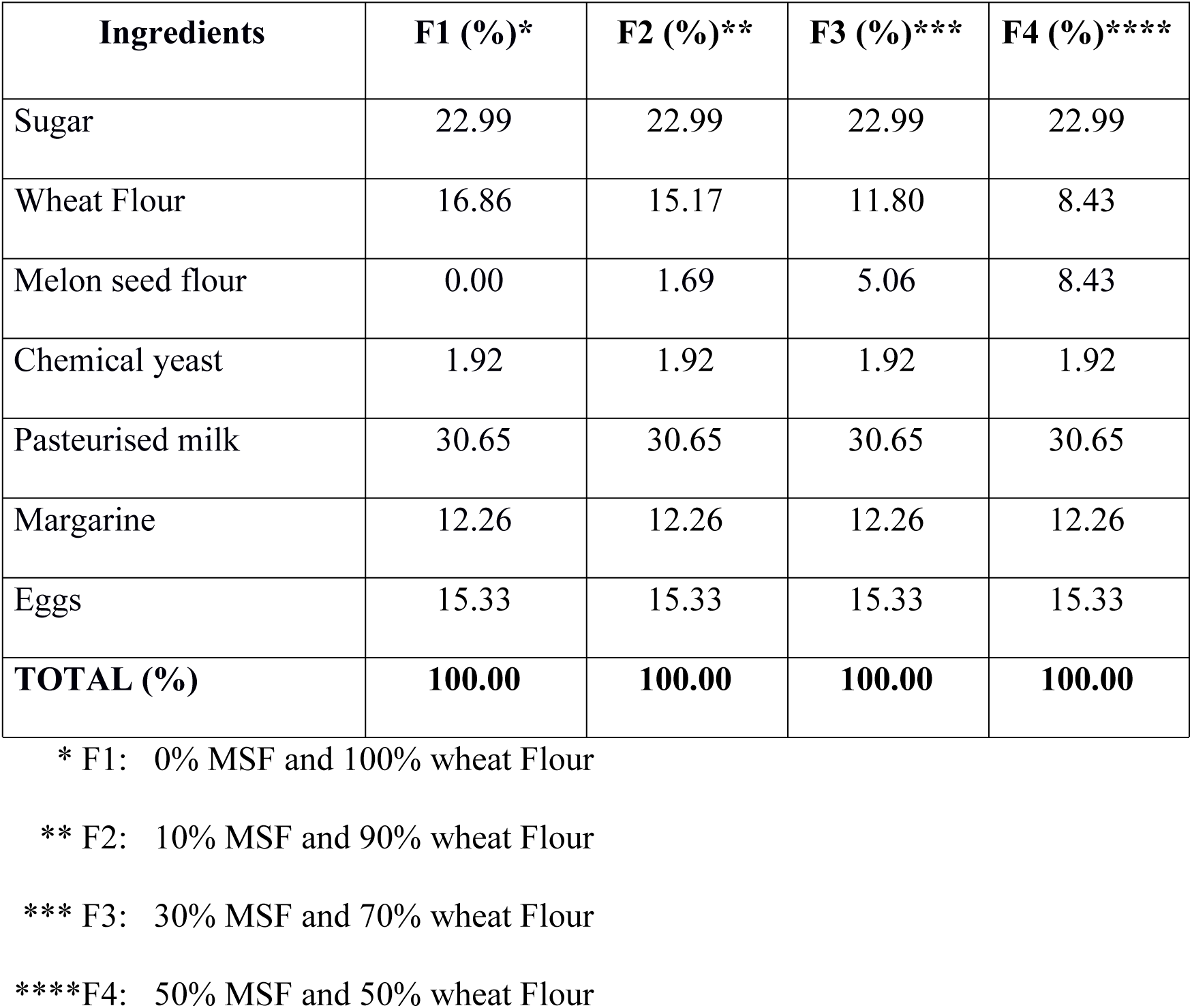
Formulations of cakes containing Cantaloupe (*Cucumis melo* L. var. *reticulatus*) Melon seed flour.

For cake development, all the ingredients were beaten for 15 minutes in a mixer until a homogeneous mass was obtained. Subsequently, the dough was placed in aluminium roasters (45 cm × 30 cm dimensions) and placed in a preheated baking oven (180°C average temperature). The heat treatment ensured that the geometric centre of the food reached temperatures above 70°C. The cakes were removed from the oven after checking their cooking and cooled on a bench at room temperature.

All formulations were processed under the same conditions, utilising the same equipment, and preparation procedures. The microbiological standard was assured by means of microbiological analysis, as described in Brazilian law [26].

#### 2.6.3 Global Acceptance

The four formulations were evaluated for Global Acceptance, using a nine-point structured hedonic scale, where 9 represented the maximum grade “I liked very much”, and 1, the minimum grade “I disliked very much” [24]. In the survey, the data from 2 panelists were excluded from Global Acceptance results, due to the lack of evaluation of some samples.

Sensory evaluation occurred in individual cabins, under natural white lighting. Each panelist was instructed to read and fill in the informed consent form, as well as the sensory evaluation sheets. In addition to evaluating the acceptance of the product, participants filled out a form with information related to their socio-demographic profile. The slices of cakes (approximately 25g) were served, monadically, in white disposable plates coded with three-digit numbers and randomly accompanied by water glasses, water, and salt biscuits [24].

#### 2.6.4 *Just About Right* (JAR)

The attributes of flavour (sweetness) and texture (softness) – characteristics most cited in the literature about sensory analysis tests and considered essential for the evaluation of a product such as cake were evaluated [27,28]. The ideal attributes were measured using the JAR method with a scale ranging from 1 to 3 points. Each attribute was calculated according to the distance between intensity perception and ideal level. The obtained data were evaluated as percentage of judgments. The representations in the 3-point scale pattern for the flavour and texture attributes were as follows: 1 “less intense than I like”, 2 “ideal, the way I like it”, 3 “more intense than I like”, 1” less soft than I like”; 2 “ideal, the way I like it “, and 3 “softer than I like” [25].

### 2.7 Statistical Analysis

JAR results were presented by the frequencies of attributes intensity ratings. The evaluation of data concerning the acceptance of cake with melon seed flour was carried out using the ANOVA statistical analysis followed by the Tukey average test (p<0.05). To evaluate panelists individual preferences, the Internal Preference Map was performed through Principal Component Analysis (PCA), using the XLSTAT statistical package.

## 3. RESULTS AND DISCUSSION

### 3.1 Centesimal Composition

Centesimal composition and dietary fibre of MSF are in Table 2. The values found highlight the nutritional potential of this residue, especially regarding dietary fibre, lipids, and proteins content.

**Table 2.**
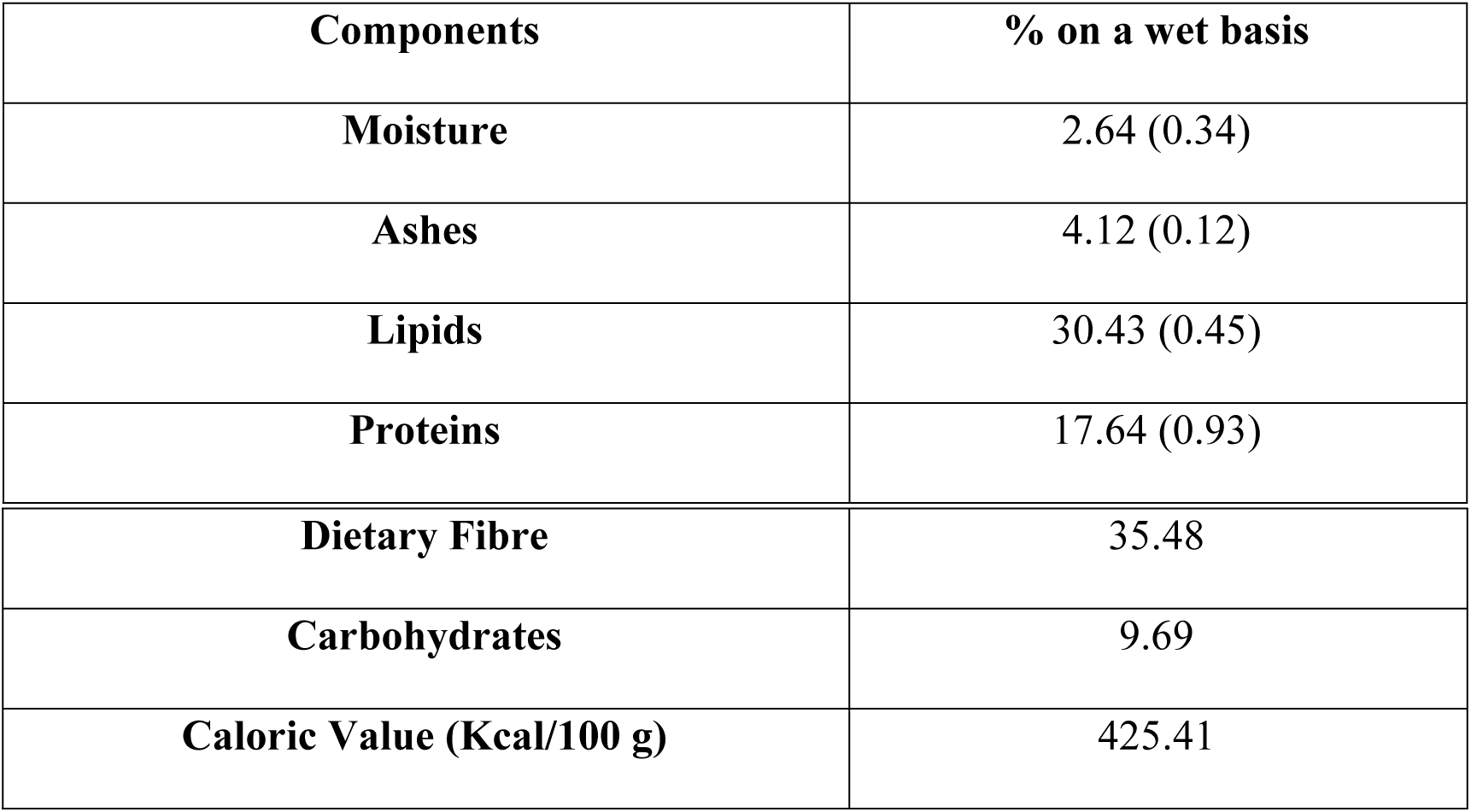
Centesimal Composition, Dietary Fibre, and Energy Value of Cantaloupe (*Cucumis melo* L. var. *reticulatus*) Melon seed flour.

| Components | % on a wet basis |
| --- | --- |
| Moisture | 2.64 (0.34) |
| Ashes | 4.12 (0.12) |
| Lipids | 30.43 (0.45) |
| Proteins | 17.64 (0.93) |
| **Dietary Fibre** | 35.48 |
| **Carbohydrates** | 9.69 |
| **Caloric Value (Kcal/100 g)** | 425.41 |

According to Table 2, the moisture content in MSF was low (2.64%), which is a positive contribution to maintaining organoleptic characteristics and microbiological quality, in addition to meeting the specific requirements for flours, cereal starch, and sharps, which establish a maximum quantity equivalent to 15.0% [29]. Moreover, the fact that the seeds have inherently low humidity, which is associated with the drying process to which the seeds were submitted, allows the concentration and preservation of the nutrients, thus decreasing perishability and promoting sustainable utilisation [30,31].

Moisture, ashes, and carbohydrates results are similar to those obtained in studies by Petkova and Antova [32], in which the authors investigated the centesimal composition of seeds of three different species of *Cucumis melo L*. melon from Bulgaria (*Honeydew Dessert* 5 and *Hybrid*1), as well as the study by Umar et al. [16] which analysed the physic-chemical composition of *Citrullus ecirrhosus* melon seeds.

In general, based on the results obtained by the centesimal composition and dietary fibre analysis, the raw material extracted from melon seeds are an ideal alternative source of functional food to promote health due to the high content of protein (17.64), lipids (30.43), and fibre (30.43). Mehra et al. [33] analysed dry seeds of *Cucumis melo* melon and found lipid values (37.17%) similar to the ones obtained by this research, although the dietary fibre values were lower. In a review study, Silva et al. [6] showed, through different works with *Cucumis melo* L. melon seeds, that the values of protein oscillated from 15% to 36%. In the work of Moura Rolim et al. [19], cantaloupe melon seeds presented the following nutrient contents in 100g of sample: 3.1g of moisture; 22.06 g protein; 24.56 g of lipids and 15.4 g of total fiber.

The high content of lipids occurring in MSF is beneficial because this reduces the concentration of fats added to products like cakes, biscuits, breads, etc. This is justified because, for the preparation of the cakes of the aforementioned study, the tests confirmed the possibility of reducing the margarine *per capita* used in the preparation of the formulations.

As a result of the present research, the fibre content obtained in the MSF analysis (35%) stands out. According to ANVISA [34], a product is considered as a source of dietary fibre when it has at least 6% of this element. In addition to providing an important amount of this nutrient, it is observed that the melon seed flour is also rich in lipids and contains good fractions of proteins, thus making it ideal to serve as an ingredient in the manufacture of any food.

### 3.2 Mineral Analysis

The concentrations of mineral salts present in the MSF are shown in Table 3. The occurrence of phosphorus (1508 mg/100 g), potassium (957 mg/100 g), and magnesium (504 mg/100 g) stands out. Morais et al. [35] also reported the prominence of these last two minerals in the micronutrients quantification. Accordingly, Fundo et al. [11] confirmed the highest concentrations of potassium, mainly located in the residual parts (peels and seeds), corresponding to 44% of the total content, with the seeds representing 14% of the total potassium level.

**Table 3.** Values of minerals in Cantaloupe Melon (*Cucumis melo* L. var. *reticulatus*) seed flour.

| Minerals | Values (mg/100 g) |
| --- | --- |
| Calcium | 36.54 |
| Copper | 1.61 |
| Total Chromium | 0.01 |
| Iron | 5.42 |
| Total Phosphorous | 1507.62 |
| Magnesium | 504.03 |
| Manganese | 4.43 |
| Total Nickel | 0.08 |
| Potassium | 957.35 |
| Selenium | 0.00 |
| Sodium | 18.50 |
| Zinc | 8.38 |

The intake of 100 g of MSF meets the requirements established by the Dietary References Intakes – DRI’s for minerals for men and women aged 19-70 years is as follows: Cr (35 × 10^−6^ and 25 × 10^−6^ mg/d), Mg (400-420 and 310-320 mg/d), P (700 mg/d, both genders), and Zn (11 and 8 mg/d), and should not exceed intakes of K (4,700 mg/d) and Na (1,500 mg/d).

The predominant minerals identified in this study (P, K, Mg) were similar to those obtained by Pimentel et al. [36]. Moreover, Azhari et al. [37], found corresponding results for calcium (Ca) and copper (Cu) while disclosing higher concentrations of potassium (9,548.33 mg/100 g), sodium (386.13 mg/100 g), zinc (44.03 mg/100 g), iron (81.17 mg/100 g), magnesium (3,299.27 mg/100 g), and manganese (15.20 mg/100 g) in the oil extracted from *Cucumis melo* var. *tibish* melon seeds.

It is noteworthy that the different planting conditions influence the concentration of minerals in the melon, and consequently, their distribution into the residual parts of the fruit. Melon seeds are an important source of minerals, namely phosphorus, potassium, and magnesium, which are found in abundance in MSF. As for the others, although the values are not as representative, it must be considered that this portion of the food (seeds) is the smallest fraction of the melon.

### 3.3 Fatty Acids Profile

The amount of fatty acids expressed as g/100 g of melon seed flour and as g/100 g of lipids are in Table 4. It is possible to notice that MSF has a high percentage of fat, with a high content of polyunsaturated fatty acids and a low concentration of saturated fatty acids. Altogether, the composition of mono and polyunsaturated fatty acids accounts for 80% of the total lipids in the flour. It should be noted that this flour is rich in omega-6 fatty acids (linoleic), whose concentration was 64% of the total fats in the sample, and also contain a considerable amount of oleic acid (15.9%).

**Table 4.** Values of fatty acids in the Cantaloupe Melon (*Cucumis melo* L. var. *reticulatus*) seed flour.

| Determination | g/100 g of MSF | g/100 g of lipids |
| --- | --- | --- |
| **Fatty Acids** |  |  |
| Saturated | 3.95 | 13.9 |
| Monounsaturated | 4.56 | 16.2 |
| Polyunsaturated | 18.01 | 63.8 |
| Total trans-isomers | ND < 0.01<sup>a</sup> |  |
| N.I. | 0.47 | 1.7 |
| **Fatty Acids Composition** |  |  |
| C16:0 palmitic | 2.57 | 9.1 |
| C16:1 omega 7 palmitoleic | 0.03 | 0.1 |
| C17:0 margaric | 0.02 | 0.1 |
| C 18:0 stearic | 1.29 | 4.6 |
| C 18:1 omega 9 oleic | 4.50 | 15.9 |
| C 18:2 omega 6 linoleic | 17.95 | 63.5 |
| C 20:0 arachidic | 0.04 | 0.1 |
| C 18:3 omega 3 $\alpha$ alpha linolenic | 0.06 | 0.2 |
| C 20:1 omega 11cis-11eicosenoic | 0.03 | 0.1 |
| N.I. | 0.47 | 1.7 |
| C 24:0 lignoceric | 0.03 | 0.1 |

<sup>a</sup>ND = Not detected / NI = Not identified.

Similar results were found by Bouazzaoui et al. [14], Azhari et al. [37], and Mallek-Ayadi et al. [38], who identified the linoleic as the main polyunsaturated fatty acid, with respective values to 60%, 61% and 69%, followed by oleic acid (25%, 19, and 16%), as well as the saturated fatty acids, palmitic (10%, 10%, and 9%) and stearic (5%, 9%, and 6%). Silva et al. [2], confirmed the high levels of linoleic acid (ranging from 52 to 69%) in several studies, which is in agreement with the values found for the MSF.

It has been observed that MSF contains a significant amount of mono and polyunsaturated fatty acids, whose recommendation is consonant for most individuals. MSF is rich in omega-6 polyunsaturated fatty acid (17.95 g /100 g) and has a considerable value of monounsaturated omega-9 fatty acid (4.50 g/100 g). These characteristics can configure a protective cardiovascular effect, since these polyunsaturated and monounsaturated fatty acids contribute to the reduction of LDL-cholesterol and total cholesterol, in addition to their antioxidant properties stemming from the presence of carotenoids and vitamin E. They can also act in the prevention of diseases, such as cancer [39-41].

### 3.4 Morphological Analysis

The micrographs produced by SEM revealed some spherical particles of different diameters (Figure 1), most of them with smooth surface (Figures 1d, 1e, 1f), homogeneous size, granular appearance, and formation of structural groupings. The presence of hemicellulose and lignin (main components) was associated with the presence of dietary fibres and constituents of cell wall (cellulose) (Figure 1b) [42, 19].

**Figure 1.**
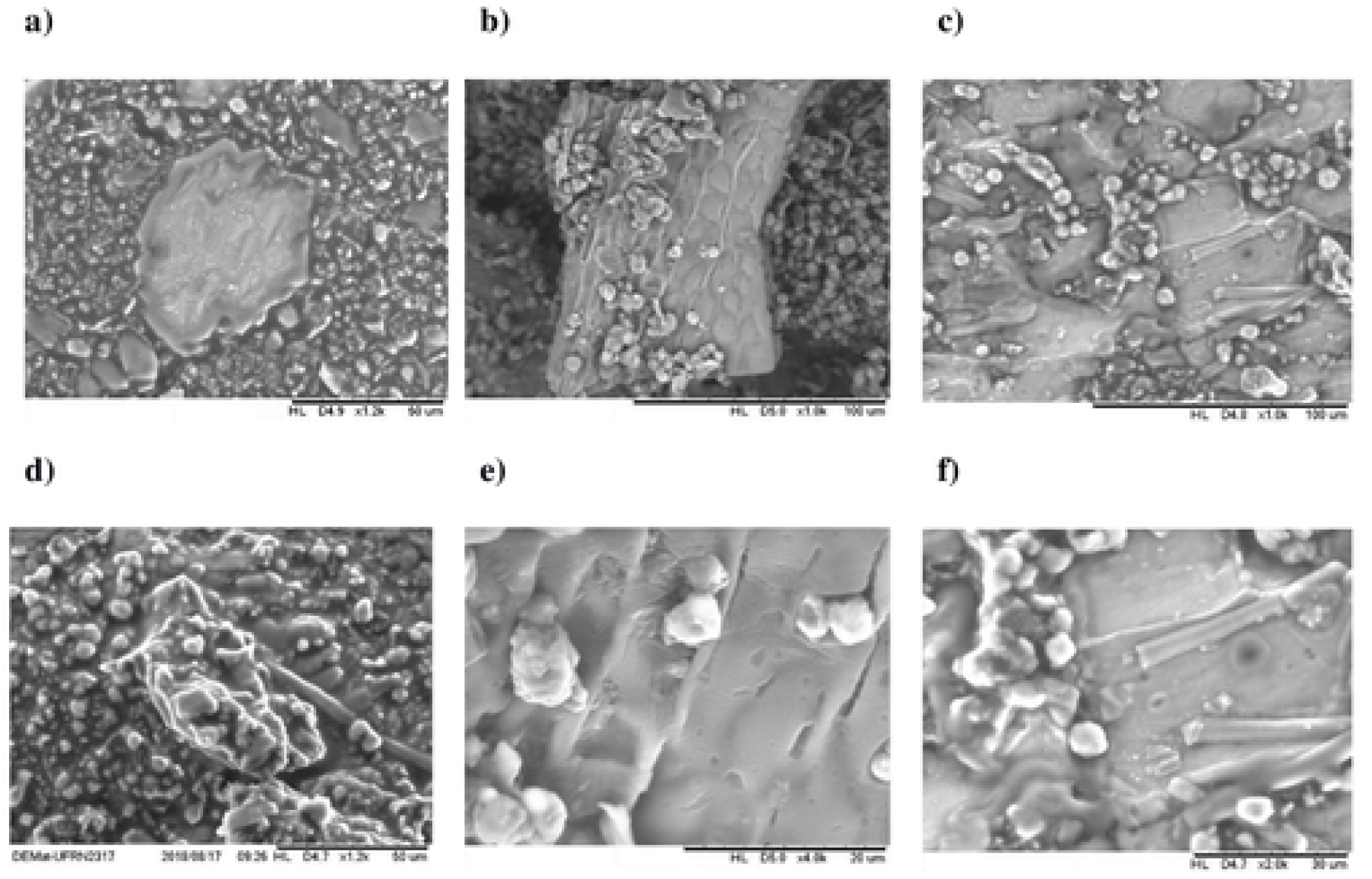
Scanning electron microscopy of Cantaloupe Melon (*Cucumis melo L.* var. *reticulatus*) seed flour: (a) 1,2 k, (b) 1,0 k, (c) 1,0 k, (d) 1,2 k, and 4,0 k, and (f) 2,0 k. Figure 1e represents two configurations; the heaviest particle indicates a higher atomic number; therefore, it has a stronger glow (clearest picture). In contrast, the lighter elements (lower atomic numbers), stand out because they are darker.

The melon seed is composed of three clearly differentiated parts: tegument, endosperm cells (middle seed layer) and perisperm (inner seed layer). The integument can be defined as the wall of the seed, also called the endocarp [38]. The resistance to digestion, even after the treatment methods, indicated an external layer consisting of fibrous, mostly insoluble, filaments (Figure 1c, 1d, 1f), protecting the embryo and the endosperm (Figures 1a, 1b) to allow for the storage of proteins and lipids [19]. The representation of a structured arrangement with a strong tegumentary presence reflects the permanence of seed peel remnants, reaffirming the high concentration of dietary fibre [43]. The starch is located in the perisperm and usually has granular appearance with various shapes and sizes [44]. Minerals such as phosphorus, potassium, and magnesium are likely also located in the endosperm and in globular crystals of the embryonic protein bodies. Likewise, there are also lipids and protein reserves [38].

Integrating the morphological study of MSF, the microscopic analysis, by bench magnifying glass, revealed the flour grain size, and allowed noticing its tendency of agglutination. The granules dimension ranged from approximately 1.5 to 2.0 mm, based on the microscopically obtained 10 mm diameter (Figure 2).

**Figure 2.**
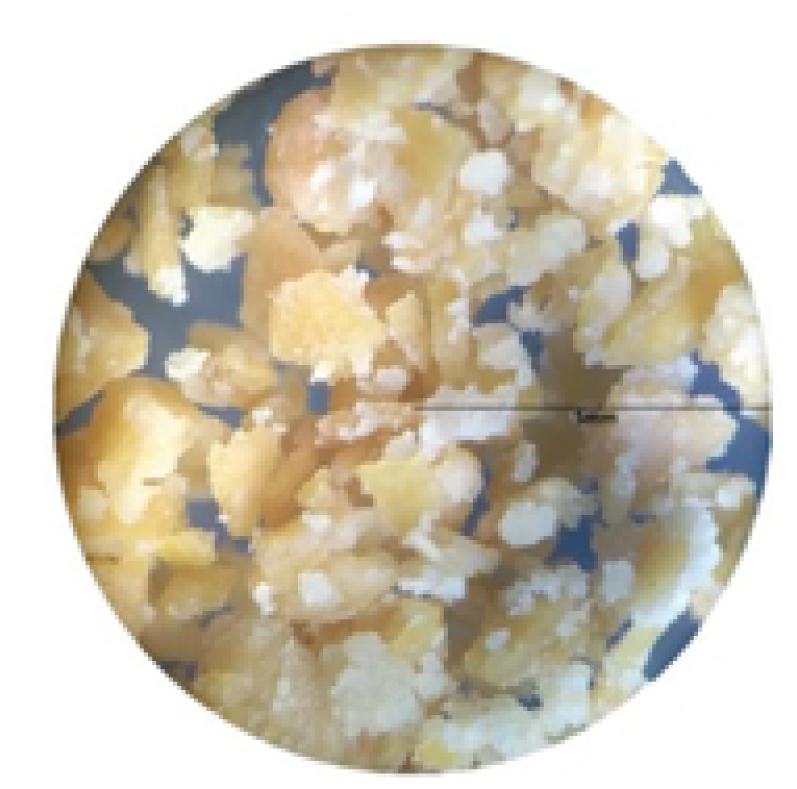
Image of Cantaloupe Melon (Cucumis melo *L. var*. reticulatus) seed flour granules, dimensioned by bench magnifying glass.

### 3.5 Sensory Analysis of Cake Formulations

Prior to the Sensory Analysis, all cake formulations and the melon seed flour were submitted to microbiological analyses, which generated favourable results for product consumption, since their sanitary and hygienic conditions were guaranteed.

#### 3.5.1 Global Acceptance

Upon analysis of the Global Acceptance results, the only statistically significant difference occurred between the formulations F2 (containing 10% MSF) and F4 (with 50% MSF). Formulations F1 (standard – 0% MSF) and F3 (30% MSF) were like the others with no significant differences, including F4 (the formulation containing the higher amount of MSF). It should be noted that F2 exhibited the highest acceptance, although this formulation was statistically equal to F1 and F3, with F1 and F3 being statistically equivalent to each other and to F4 (Table 5).

**Table 5.** Global Acceptance Results for cakes prepared with different contents of Cantaloupe Melon (*Cucumis melo* L. var. *reticulatus*) seed flour.

| FORMULATIONS | GLOBAL ACCEPTANCE |
| --- | --- |
| F1 | 6.56<sup>ab</sup>(±1.95) |
| F2 | 7.08<sup>a</sup> (±1.61) |
| F3 | 6.59<sup>ab</sup> (±1.83) |
| F4 | 6.10<sup>b</sup> (±2.17) |

<sup>ab</sup> Means followed by the same lowercase letters do not differ by the Tukey test (p>0,05).
F1 – 0% MSF and 100% wheat meal.
F2 – 10% MSF and 90% wheat meal
F3 – 30% MSF and 70% wheat meal
F4 – 50% MSF and 50% wheat meal

For each formulation, most panelists assigned grade 6 (“slightly liked”) to the formulation that contained 50% MSF, grade 7 (“I liked regularly”) to the cake containing 30% MSF, grades 8 (“I liked it very much”) and 9 (“I liked it tremendously”) for the product containing 10% of melon seed flour. Upon analysing these results, we observed that cake F2 was the one that pleased the evaluators.

In general, the high acceptability of the cakes produced with MSF is confirmed, with the formulation prepared exclusively with wheat flour showing lower acceptance when compared to products containing 10% and 30% concentrations. Moreover, even with the formulation containing 50% of melon seed flour performing the worst in terms of taste, this result was similar to those referring to the other formulations.

In order to consider the individual responses of each taster, not just the group average, the Principal Component Analysis (PCA) was applied, using a four samples × 133 taster’s dimension matrix. Data were adjusted and the acceptance test results were presented by a Bi-plot graph (internal preference vector map) as shown in Figure 3. The map represents the panelists as vectors and the samples dispersed among them.

**Figure 3.**
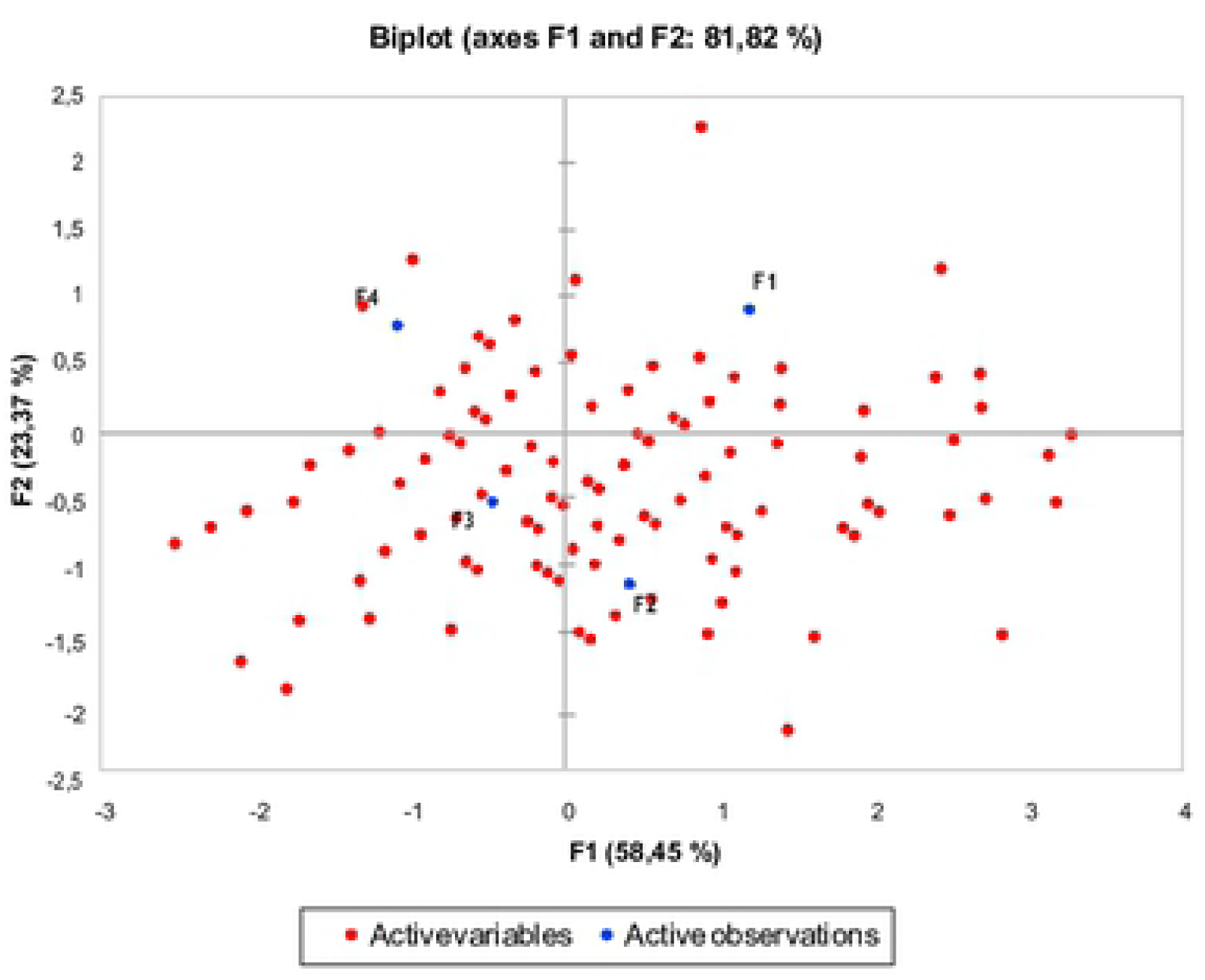
Internal Preference Vector Map for 4 cake formulations: F1 – 0% MSF and 100% wheat flour, F2 – 10% MSF and 90% wheat flour, F3 – 30% MSF and 70% wheat flour; F4 – 50% MSF and 50% wheat flour, evaluated in accordance with the 9-point Hedonic Scale.

In order to ensure success of the PCA methodology, it is ideal that two components accumulate a percentage of variance equal to or greater than 70% [45]. The variability analysis demonstrated that the composition of two components explained 81.82% of the variance. In the graph (Figure 4), all formulations are distributed in an equivalent manner without dispersion of panelists for a specific sample, demonstrating that all the cakes were well accepted. Notably, PCA had a slightly higher vector density involving F2 (which contains 10% MSF). This indicates a consumer preference for this formulation, as 75% of the grades given by the judges in formulation 2 corresponded to values between 7 and 9 for overall acceptance. Regarding F4 (cake made with 50% MSF), we concluded that this formulation was the least accepted, as F2 and F4 were in opposite quadrants. In contrast, the other formulations did not differ statistically with respect to the standard formulation. Therefore, PCA showed that the consumers evaluated the samples in an analogous way, and we conclude that the addition of MSF to the standard cake did not influence the preference of the taster.

**Figure 4.**
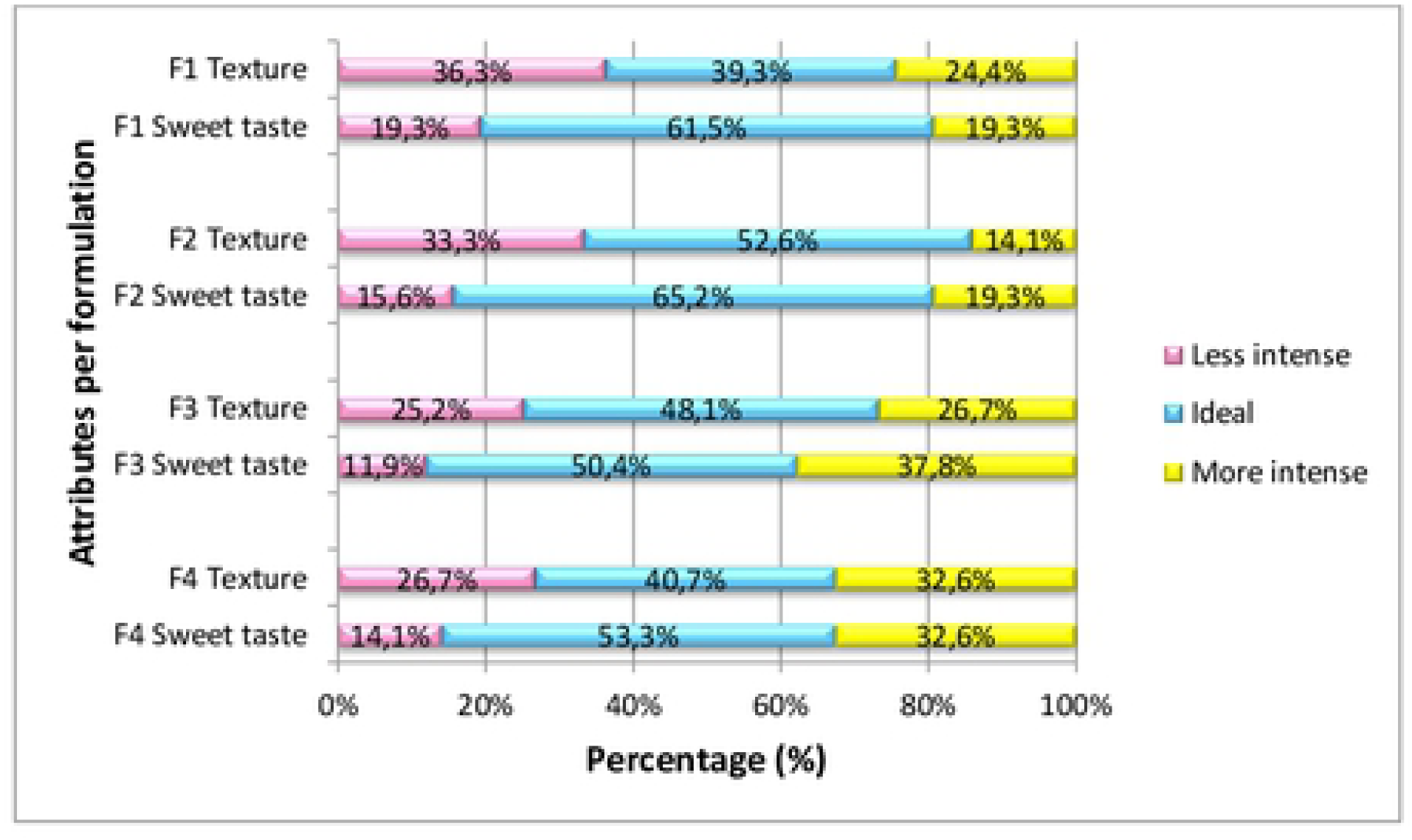
Frequencies of intensity ratings (1 = “less intense than I like”, 2 = “ideal, the way I like it”, and 3 = “more intense than I like”) for the sweet taste attribute and (1 = “less soft than I like”, 2 = “ideal, the way I like it”, 3 = “softer than I like”) for the attribute texture. Measured on a 3 – point JAR Scale for different cakes formulations (F1 – 0% MSF and 100% wheat flour, F2 – 10% MSF and 90% wheat flour, F3 – 30% MSF and 70% wheat flour, and F4 – 50% MSF and 50% wheat flour).

As well as highlight the benefit of using MSF, creating a product with considerable energetic value, source of proteins and fibres, and serving as an alternative for the enriching of preparations, Abelama et al. [46] determined the potential use of jackfruit seed flour for the preparation of biscuits, breads, and cakes.

Oliveira et al. [47] conducted a study investigating the partial substitution of corn granules by grape waste flour (peels and seeds) using the 9-point Hedonic Scale for checking the acceptance, and discovered a better acceptance for the formulations containing, 15% and 20% (in order of preference), which also offered the highest fibre content. Another investigation of grape waste flour (peels) showed that the Global Acceptance Test disclosed that the extrudate containing 5% of this flour had better acceptance than the control formulation and the 10% formulation (regarding the texture parameter), and was not different from the control formulation in terms of flavour [48].

It is worth noting the good acceptance of the 10% and 30% MSF cake formulations, which is in line with the excellent results demonstrated in the mentioned research. In addition, the greater addition of flour to the cake (formulation containing 50% MSF) implied a slight reduction in the acceptance percentage. The results of this study are similar to that of Storck et al. [49], involving the sensorial analysis of cakes, among them that of melon seeds, which was considered one of the preparations that presented higher fiber content, besides acceptance of global characteristics.

#### 3.5.2 *Just About Right* (JAR)

The JAR scale is used in different types of evaluations, both food-related and others, since its objective is to define the most appropriate attribute (ideal level) for a specific food, in addition to being straight-forward for non-trained evaluators to establish their preferences [50].

The sensory analysis participants evaluated the cakes containing 0%, 10%, 30%, and 50% of MSF for texture (softness) and flavour (sweetness). Observing Figure 5, we can see that the panelists preferred cake F2 and consider the texture (52.6%) and flavour (65.2%) of this formulation as “ideal”.

Evaluating the attributes separately, the preference for texture was equivalent for formulations F2 and F3, as well as for the formulations F1 and F4, with the values being higher for the first group. Regarding the sweet taste attribute, F1 and F2 were considered equivalent, as well as F3 and F4, highlighting the first two formulations mentioned as the preferred ones of the consumers

In order of choice, the panelists preferred samples F2 (53%), F3 (49%) and F4 (41%) (in that order) for the texture attribute (softness), which contained 10%, 30%, and 50% of MSF, respectively. Their last option was formulation 1, which had no addition of MSF (corresponding to 39% on the ideal scale).

For the sweetness attribute, the evaluators presented the following cake preference sequence: F2 (65%), F1 (62%), and F4 (53%), containing 10%, 0%, and 50% of MSF, respectively. The F3 cake (with 30% MSF) was the least preferred, representing 50% preference on the ideal scale. These results highlight the disposition to choose products that are similar to the traditional ones (refined).

Although some panelists attributed grades 1 and 3 for certain analysed formulations, grade 2 (representative of “ideal” flavour and texture) was predominant among the responses for all cake samples for these two attributes. This indicates that although most consumers are not accustomed to whole food intake, these products have the potential to be well accepted if they are introduced to the eating habits of individuals after nutritional education work.

We observed a higher preference for cakes without MSF and for the one with the lowest MSF concentration (in flavour evaluation), as shown in Figure 4. These findings are similar to those observed in Guimarães, Freitas, Silva [51] in a study using watermelon bark seam flour in the preparation of cakes.

Similar to the present study, work developed by Spada et al. [52], which used different formulations (50%, 75% and 100%) of jackfruit flours to substitute cocoa powder for the preparation of cappuccinos, observed that the lowest concentration ratios used (50% and 75%) in the research, obtained the best results of acceptance in the sensorial analysis, still perfecting some characteristics of the original product.

In general, all MSF containing cakes formulations were well accepted, even with increasing concentrations of this ingredient. It is noteworthy that, especially for the texture attribute, the cake that was the least well received by the panelists was the one without addition of MSF. Conrad et al. [53], studying the properties of melon seed flour, *maazoun* variety, confirmed that the emulsifying ability of this product presented better results than wheat flour, confirming that melon seed meal can potentially be added to products of baking.

Comparing nutritional value of a traditional cake produced exclusively with wheat flour and the cake produced with 10% MSF, we can confirm the high nutritional value of the cake added with MSF. Although the corresponding values of moisture, ash, carbohydrates, proteins, lipids, energy, sodium, and calcium were similar, there is a discrete but important increase in nutrients such as dietary fibre (0.50g/80g), magnesium (9.72mg/80g), phosphorus (37.13mg/80g), iron (0.29mg/80g), potassium (55.41mg/80g), copper (0.04mg/80g), and zinc (0.29mg/80g). In addition, considering the other percentages of MSF added to the cakes (30% and 50%), the nutritional increment is even more pronounced, contributing substantially to consumer nutrition [54].

For Rolim, Seabra and Macedo [55], a way of avoiding waste is to take advantage of all parts of the food, a process that is still little discussed and applied, since there are few studies working with this theme. Thus, it is interesting to produce information on the nutritional enrichment that these residues provide to food. In addition to minimizing losses, it increases the preparation of new preparations, adding nutritional value and enhancing the flavor, texture, aroma and color of food [56]

In this sense, the main obstacle to the use of food waste in the development of new products is the lack of awareness, training and encouragement to the food handlers to participate in the recovery initiative; presence of more conscious consumers; as well as from committed policymakers who are willing to implement consolidated waste prevention measures, allocating food adequately and sustainably [55, 57].

Thus, in addition to being nourishing, MSF is an alternative source of nutrients such as proteins, fibers, minerals, mono and polyunsaturated fatty acids, and can be an example to encourage healthy eating habits associated with sustainability concerns.

## 6. CONCLUSION

This study demonstrated the feasibility of the use of melon wastes in cake production. Given the nutritional properties found in melon seeds and the sensory acceptance of the cake produced using melon seed flour as an ingredient, is possible to propose the use of melon seed flour in foodservices to offer economic, social, and health benefits, promoting a clean production and food security by preserving food nutrients.

## ACKNOWLEDGEMENTS

The authors are grateful to the Research Support Foundation of Rio Grande do Norte – FAPERN/CAPES, Brazil – 006/2014 and to the Coordination for the Improvement of Higher Education Personnel – Brazil (CAPES), responsible for financing part of the study – Financial Code 001.

## Supporting information

**Data. The raw data for Centesimal Composition and Sensory Analysis.**

